# Loss of DAXX induces alternative lengthening of telomeres (ALT)-associated hallmarks in prostate cancer cells in a context-specific manner

**DOI:** 10.1101/2025.08.06.668744

**Authors:** Joakin O. Mori, Joseph Da, Jiyoung Kim, Anthony Rizzo, Christine Davis, Carlo Lanza, Jacqueline A. Brosnan-Cashman, Alan K. Meeker, Christopher M. Heaphy, Mindy K. Graham

## Abstract

**Background:** The histone chaperone complex, consisting of the death domain–associated protein (DAXX) and the alpha-thalassemia/mental retardation X-linked protein (ATRX), plays a pivotal role in maintaining chromatin through the deposition of the histone variant H3.3. Mutations leading to loss of ATRX or DAXX function are linked to the non-telomerase, alternative lengthening of telomeres (ALT) phenotype in certain cancers. Engineered ATRX mutations have previously been found to induce features of ALT in prostate cancer cell lines, notably in LAPC-4, but not in CWR22Rv1. This study determined the impact of *DAXX* mutations on ALT-associated characteristics in CWR22Rv1 and LAPC-4.

**Methodology:** Mutations were induced in CWR22Rv1 and LAPC-4 cells by targeting exon 2 of *DAXX* using the CRISPR-Cas9 genome editing strategy. The resulting mutant clones were then evaluated for ALT-associated characteristics, including the presence of ALT-associated PML bodies (APBs), C-circles, telomere length heterogeneity, and a lack of telomerase activity.

**Results:** Four CWR22Rv1 *DAXX* mutant clones (*DAXX*^*Mut*^*1*-*4*) and five LAPC-4 clones (*DAXX*^*Mut*^*1-5*) were evaluated. In CWR22Rv1, *DAXX*^*Mut*^*1, DAXX*^*Mut*^2, and *DAXX*^*Mut*^*4* were true knockout clones with frameshift mutations in both copies, while CWR22Rv1 *DAXX*^*Mut*^*3* had a frameshift mutation in one copy and an in-frame mutation in the other. Protein expression was undetectable in all the CWR22Rv1 clones, including CWR22Rv1 *DAXX*^*Mut*^*3*. In LAPC-4, *DAXX*^*Mut*^*1* was a true knockout, while *DAXX*^*Mut*^*2, DAXX*^*Mut*^*3, DAXX*^*Mut*^*4*, and *DAXX*^*Mut*^*5* clones had at least one in-frame mutation. Among these LAPC-4 clones, only *DAXX*^*Mut*^*1* had undetectable protein by western blotting. ALT-associated characteristics such as APBs, C-circles, and telomere length heterogeneity were observed only in CWR22Rv1 *DAXX*^*Mut*^*4*. All the clones maintained telomerase activity, regardless of whether ALT-associated hallmarks were observed.

**Implications:** The CWR22Rv1 and LAPC-4 DAXX mutant clone models provide useful tools for future studies on telomere maintenance mechanisms and DAXX-related biology, particularly in prostate cancer.

## INTRODUCTION

Chromatin maintenance at repetitive DNA sequence regions (including centromeres, pericentromeres, and telomeres) of the genome is critical to prevent the accumulation of chromosomal instability. Telomeres, repetitive hexameric sequences bound by the protein complex shelterin and located at the ends of eukaryotic chromosomes, maintain genomic stability and integrity by preventing recognition of the chromosomal ends as double-stranded breaks and exonucleolytic degradation [1]. Telomerase is a telomere-specific reverse transcriptase responsible for maintaining telomere length. Telomerase activity varies in cells depending on a cell’s self-renewal and replicative capacity. Embryonic stem cells, which can self-renew and replicate indefinitely in culture, maintain high telomerase activity compared to adult stem cells that have reduced self-renewal and replicative capacity [2-5]. Telomerase activity is absent in normal somatic cells [4]. Consequently, telomeres progressively shorten in somatic cells with each round of cell division because of incomplete DNA replication in the lagging strand [6]. Critically short telomeres induce p53-dependent replicative senescence or apoptosis in normal cells [7]. Most cancer cells reactivate telomerase, thereby circumventing telomere shortening-induced replicative senescence or apoptosis [8 9]. In rare circumstances, cancer cells circumvent critically short telomere-induced replicative senescence or apoptosis by activating telomere-lengthening mechanisms [10]. Most cancers upregulate telomerase; however, 10-15% of cancers lack telomerase activity [5 11]. A subset of telomerase-negative cancers maintain their telomere lengths by a telomere-specific mechanism of homology-directed repair called Alternative Lengthening of Telomeres (ALT) [12-14].

The histone chaperone alpha-thalassemia/mental retardation X-linked protein (ATRX) and death domain–associated protein (DAXX) complex is involved in the maintenance of chromatin through the deposition of the histone variant H3.3 [15 16]. Previously, we and others observed a robust correlation between ALT and mutations found in the *DAXX* and *ATRX* genes across pancreatic neuroendocrine tumors (PanNETs), pediatric glioblastoma, and uterine leiomyosarcomas [17-19].

The *DAXX* gene, located on chromosome 6, encodes a multifunctional protein thought to reside primarily in the nucleus [20], although it has been previously shown to be capable of translocating to the cytoplasm [21 22]. In the nucleus, the DAXX protein is known to have roles in chromatin maintenance and transcriptional regulation. While the DAXX protein works in combination with ATRX to prevent the accumulation of double-strand breaks at telomeric DNA, it also acts independently of ATRX to prevent the accumulation of double-strand breaks in centromeric DNA [23].

We have previously demonstrated that the introduction of *ATRX* loss-of-function mutations induced ALT features in LAPC-4 prostate cancer cells but not in CWR22Rv1 prostate cancer cells, suggesting induction of ALT-associated features following ATRX loss is context-specific [24]. As it relates to telomere regulation in prostate cancer, the role of DAXX is not well understood. However, previously published studies suggest interactions with multiple key players of prostate carcinogenesis. Androgen receptor (AR) transcriptional activity is critical for the growth and survival for the majority of prostate cancers. Interestingly, DAXX has also been reported to bind to the AR in prostate cancer cells, preventing ligand-dependent transactivation and binding to androgen response elements in the genome, which consequently downregulates AR transcriptional activity [25]. The DAXX protein is also thought to regulate the subcellular localization of PTEN, a tumor suppressor protein, although the molecular mechanism underlying this regulatory role has not been well elucidated [26]. Notably, PTEN is frequently mutated in prostate cancer, with roughly half of treatment-resistant prostate cancers having PTEN loss [27]. The regulatory role of DAXX on AR and PTEN activity suggests an antitumor role. However, DAXX has also been shown to exhibit a protumorigenic role in prostate cancer [28 29]. DAXX expression was shown to be associated with increased odds of *TMPRSS2-ERG* gene fusions and *ERG* expression, advanced Gleason grade and pathological staging, and increased risk of biochemical recurrence [28]. *TMPRSS2-ERG* fusion is a common event in prostate cancer – present in approximately 50% of cases. *ERG* is typically not expressed in normal prostate. However, the fusion of the *TMPRSS2* promoter with the *ERG* gene results in the abnormal expression of *ERG* in an AR-dependent manner. Separately, DAXX can promote prostate tumorigenesis by transcriptionally suppressing autophagy, a process associated with tumor suppression [30 31]. Specifically, DAXX has been shown to repress the expression of autophagy modulator genes *DAPK3* and *ULK1* [30] by binding and recruiting DNA methyltransferase 1 (DNMT1) to the enhancer or promoter regions [31].

In this study, we investigated ALT-associated molecular and cellular characteristics in prostate cancer cell lines, CWR22Rv1 and LAPC-4, after introducing *DAXX* mutations using CRISPR-Cas9-mediated genome editing. CWR22Rv1, originally derived from a primary tumor [32], and LAPC-4, derived from lymph node metastasis [33], are telomerase-positive prostate cancer cell lines. We introduced mutations in the second exon of *DAXX*, which encodes the DAXX Helix Domain that interacts with ATRX [34]. Multiple clones were generated harboring in-frame and out-of-frame mutations, which were confirmed by Sanger sequencing. In a context-dependent manner, mutations in *DAXX* were associated with loss of protein expression and induced some ALT-associated features, including ALT-associated PML bodies (APBs) and ALT-associated C-circles, but not telomere length heterogeneity.

## MATERIALS AND METHODS

### Cell culture

CWR22Rv1 was cultured in RPMI 1640 medium (Gibco) supplemented with 10% heat-inactivated fetal bovine serum (FBS, Sigma) and 1% mixture of Penicillin 10,000 units/mL and Streptomycin 10,000 μg/mL (Pen/Strep, Quality Biological). LAPC-4 was cultured in IMDM medium (Gibco) supplemented with 10% FBS, 1% Pen/Strep, and 1 nM of R1881. U2OS was cultured in DMEM (Gibco) supplemented with 10% FBS and 1% Pen/Strep. All cell lines were submitted to the Genetic Resources Core Facility at Johns Hopkins for mycoplasma detection and cell line authentication by short tandem repeat profiling (Promega).

### Inducing DAXX knockout mutations

*DAXX* knockout mutations were introduced using a previously described CRISPR-Cas9 genome editing strategy [23 24]. Briefly, two CRISPR-Cas9 nickase guide RNAs (gRNA-1: 5’-CATGAGGCTCAGAGGAGCTA, gRNA-2: 5’-AGAGGAAGCAGTAGTTCGGG) were designed to target exon 2 of *DAXX* using CRISPR Design (MIT CRISPR tool [35]). The gRNAs were cloned into the GFP-expressing Cas9n plasmid, PX461, a gift from Feng Zhang (Addgene #48140). Lipofectamine 3000 (Thermo Fisher Scientific) was used to transfect either empty vector PX461 or co-transfect both DAXX gRNA1-PX461 and DAXX gRNA 2-PX461 into CWR22Rv1 and LAPC-4 cells. FACS-sorted GFP-positive cells were plated in 150 mm dishes. Cell colonies were isolated using cloning cylinders (Sigma-Aldrich) and screened for DAXX protein by immunostaining. Promising knockout clones were subsequently validated by western blotting and Sanger sequencing.

### Western blotting

Western blot was performed as described previously [24]. Frozen cell pellets were resuspended in 100 mL of RIPA buffer (Cell Signaling) and incubated on ice for 30 minutes. Cell lysates were centrifuged for 30 minutes at 13,000 RPM at 4°C. Supernatant was collected, and the BCA Protein Assay Kit (Pierce) was used for protein concentration. An equal volume of 2X Laemmli Buffer (Bio-Rad) containing 5% 2-mercaptoethanol (MilliporeSigma) was added to the cell lysate and incubated for 10 minutes at 95°C. Denatured lysate was cooled on ice and 30 mg of cell lysate was loaded into the well of NuPAGE™ 3-8% Tris-Acetate Protein Gel (Invitrogen). Samples were run for 1-1.5 hours at 150 volts in NuPAGE™ tris-acetate SDS running buffer (Invitrogen). Electrophoretically separated proteins were transferred to a nitrocellulose membrane (Bio-Rad) in Tris-Glycine transfer buffer (Bio-Rad) containing 5% methanol (MilliporeSigma) for 3 hours at 325 mA at 4°C. Membranes were blocked in 5% milk (Bio-Rad) in tris buffered saline with Tween 20 (TBST, MilliporeSigma) for one hour, and then incubated in primary antibody overnight at 4°C in 5% milk in TBST. The following dilutions in 5% milk were prepared for each primary antibody: 1:20,000 GAPDH (Cat# D16H11, Cell Signaling), 1:10,000 beta-actin (Cat# 8H10D10, Cell Signaling), or 1:2,000 DAXX (Cat# HPA008736, Sigma). Membranes were washed in TBST and incubated in secondary antibody for one hour in 5% milk in TBST, either 1:10,000 HRP-linked anti-rabbit IgG (Cat# 7074, Cell Signaling) or 1:20,000 HRP-linked anti-mouse IgG (Cat# 7076, Cell Signaling). Signals were detected using Clarity ECL (Bio-Rad). Blots were either exposed to film or developed using the ChemiDoc Imaging System (Bio-Rad).

### Detection of ALT-associated PML bodies

Formalin-fixed paraffin-embedded (FFPE) cell pellet blocks were prepared by fixing cell pellets in 10% phosphate-buffered formalin (MilliporeSigma) for 48 hours and embedding them in paraffin. FFPE blocks were cut into 5 µm sections for telomere fluorescence in situ hybridization (FISH) and PML immunofluorescence staining.

#### Telomere FISH

The 5 µm section slides were deparaffinized, rehydrated, incubated in 1% Tween 20 for 1 minute, and steamed for 30 minutes in citrate buffer, pH 6 (Vector Laboratories). Slides were incubated with 333 ng/mL of telomere probe, a Cy3-labeled peptide nucleic acid (PNA) oligonucleotide (N-CCCTAACCCTAACCCTAA-C) in PNA buffer (10 mM Tris pH 7.5, 70% formamide) for 5 minutes at 84°C, followed by an incubation for 2 hours or overnight at room temperature. The excess probe was washed off at room temperature with PNA buffer.

#### PML immunofluorescence staining

Slides were blocked for 10 minutes in serum-free protein block (Agilent Dako), washed in phosphate-buffered saline with Tween 20 (PBST), and subsequently incubated for two hours in 1:100 PML antibody (A301-167A, Bethyl) diluted in antibody diluent buffer (Ventana). The excess antibody was washed in PBST, and the slides were stained with anti-rabbit Alexa Fluor 647 (Invitrogen) diluted in PBS. Slides were stained with DAPI and mounted with ProLong™ Gold Antifade reagent (Invitrogen).

#### Image acquisition and scoring

Telomere FISH and PML immunofluorescence-stained slides were scanned using the TissueFAXS Plus (Tissue Gnostics) automated microscopy workstation equipped with a Zeiss Z2 Axioimager microscope. Scanned images (400X) were exported as TIFF images using TissueQuest software (Tissue Gnostics), and manually scored for cell count and co-localization of PML with ALT-associated telomeric foci with the ImageJ/Fiji cell count tool [36].

### Detection of ALT-specific C-circles

As previously described, the C-circle dot blot assay [37] was used with some modifications to detect ALT-specific C-circles. Genomic DNA was extracted from frozen cell pellets harvested from a confluent T-75 flask using the DNeasy Blood and Tissue Kit (Qiagen). DNA less than 10 kilobases were enriched using the QiaQuick PCR Purification Kit (Qiagen). C-circle signal was amplified using rolling circle replication, combining 150 ng of DNA with 2 units of phi29 DNA polymerase (New England Biolabs) in 20 mL of C-circle buffer (1x Phi29 reaction buffer, 200 ng/mL BSA, 0.1% Tween 20, 1 mM dATP, 1 mM dGTP, 1 mM dTTP). Reaction products were blotted onto a positively-charged nylon membrane (Roche) and hybridized to Digoxigenin (DIG) conjugated telomere probe (CCCTAACCCTAACCCTAACCCTAA-DIG, Integrated DNA Technologies) in DIG Easy Hyb™ buffer (Roche) overnight at 42°C. Excess probe was washed using 2X saline-sodium citrate (SSC) buffer with 0.1% SDS at room temperature, followed by 0.2X SSC 0.1% SDS at 50°C. Membranes were blocked in 5% milk TBST for at least 30 minutes at room temperature, and subsequently incubated with 1:10,000 anti-Digoxigenin-AP, Fab fragments (Roche) for at least 30 minutes. Excess antibody was washed and amplified C-circle products were detected using a CDP-Star kit (Roche). C-circle signals were quantified by densitometry using the ChemiDoc™ Imaging System (Bio-Rad).

### Telomere repeated amplification protocol (TRAP)

A previously described protocol [38] was employed, with minor modifications, to conduct the TRAP assay. Frozen cell pellets (1×10^6^ cells) were resuspended in 400 mL of NP-40 lysis buffer and incubated on ice for 30 minutes. The equivalent of 2500 cells (1 mL of crude lysate) was combined with 49 mL of 1X TRAP buffer, 200 mM dNTPs, 2 units of DNA polymerase (Platinum™ *Taq* DNA Polymerase, Invitrogen), 340 nM TS oligonucleotide (5’-AAT CCG TCG AGC AGA GTT-3’) conjugated with Alexa Fluor 488, 170 nM ACX oligonucleotide (GCG CGG CTT ACC CTT ACC CTT ACC CTA ACC), 20 aM (2×10^−17^ M) TSNT oligonucleotide (AAT CCG TCG AGC AGA GTT AAA AGG CCG AGA AGC GAT), and 340 nM NT oligonucleotide (ATC GCT TCT CGG CCT TTT). For RNase-treated lysate, 1 mL of RNase A (Qiagen) diluted to 2.5 mg/mL was combined with 8 mL of lysate and 15 mL of nuclease-free water. The reaction was incubated at room temperature for 5 minutes and 2,500 cell equivalents was used in the TRAP reaction using the following steps: 1 cycle at 30°C for 30 minutes; 1 cycle at 94°C for 10 minutes; 26 cycles of 94°C for 30 seconds, 50°C for 30 seconds, and 72°C for 45 seconds; and then holding at 4°C. To visualize the TRAP products, 10 mL of 6X nucleic acid loading buffer (Thermo Scientific) was added to each TRAP reaction. Subsequently, an aliquot of 40 mL of the reaction was loaded into a well of a 20% Novex® TBE Gel (Invitrogen), and run for 125 minutes at 200 Volts in 0.5X TBE Buffer. TRAP products labeled with Alexa Fluor 488 were visualized using the ChemiDoc™ Imaging System (Bio-Rad).

### Telomere Restriction Fragment Southern blotting

Genomic DNA extraction and Telomere Restriction Fragment (TRF) Southern Blotting were performed as previously described with some modifications [39]. Cells were resuspended in SNET buffer (20 mM Tris pH 8.0, 5 mM EDTA, 400 mM NaCl, 1% SDS) containing 400 μg/mL of proteinase K (P8107S, NEB), and incubated at 55^°^C with gentle agitation overnight. An equal volume of phenol:chloroform:isoamyl alcohol (25:24:1, pH 8, MilliporeSigma) was added. The aqueous layer was isolated, and DNA was precipitated with an equal volume of isopropanol. The DNA pellet was washed twice with 70% ethanol and air dried. Isolated genomic DNA was digested overnight with 20 units of MseI (R0525, NEB) and 0.07 units of RNase A (Qiagen) in 1XCutSmart buffer at 37°C. Digested DNA was supplemented with 20 units of AluI (R0137, NEB) and 10 units of MboI (R0147, NEB). DNA Molecular Weight Marker II, DIG-labeled (Roche) and digested DNA (2-20 ug, empirically determined input) was combined with 6X DNA loading dye (NEB) and run on 0.7% agarose TAE gel for 12 hours at 47 V. Gel was post-stained in 0.5 ug/mL ethidium bromide, and UV treated for 60 mJ using a Stratalinker [40]. The gel was subsequently incubated for 30 minutes in Denaturation Solution (0.5 M NaOH/1.5 M NaCl, Millipore), followed by 30 minutes in Neutralization Buffer (1.5 M NaCl/0.5 M Tris-HCl pH 7.4), and 30 minutes in 20X SSC buffer. DNA was transferred to positively-charged nylon membrane (Roche) overnight using the Turboblotter transfer system (GE), and subsequently UV cross-linked using a Stratalinker. Transferred TRFs were hybridized to Digoxigenin (DIG) conjugated telomere probe (CCC TAA CCC TAA CCC TAA CCC TAA-DIG, Integrated DNA Technologies) in DIG Easy Hyb™ buffer (Roche) overnight at 42°C. Excess probe was washed using 2X saline-sodium citrate (SSC) buffer with 0.1% SDS at room temperature, followed by 0.2X SSC 0.1% SDS at 50°C. Membranes were blocked in 5% milk TBST for 30 minutes at room temperature, and subsequently incubated with 1:10,000 anti-Digoxigenin-AP, Fab fragments (Roche) for 30 minutes. Excess antibody was washed and amplified TRFs were detected using CDP-Star kit (Roche). TRF sizes were determined using the ChemiDoc™ Imaging System (Bio-Rad), and telomere length distribution was recorded based on densitometry reads with the plot profile tool on ImageJ/Fiji [36].

### Statistical Analysis

Statistical tests, including t-test, were performed in R (version 4.2.0) using the rstatix package (version 0.7.2). P-values < 0.05 were considered statistically significant.

## RESULTS

### Introducing *DAXX* mutations in prostate cancer cell lines

CWR22Rv1 [32] and LAPC-4 [33] are well-characterized human prostate cancer cell lines that express full-length DAXX protein and rely on telomerase for telomere maintenance [24]. Using the CRISPR-Cas9 nickase system, we generated multiple *DAXX* mutant clones in CWR22Rv1 and LAPC4. The aspartate-to-alanine (D10A) mutation in the catalytic domain of Cas9 results in a nickase mutant, requiring two gRNAs to generate the desired double-strand break and initiate error-prone repair, thus decreasing the incidence of unwanted mutations in non-specific targets [41]. These gRNAs were designed to target a sequence in exon 2, which encodes part of the domain of DAXX that interacts with ATRX [34 42 43] and USP7 [44] (**Fig.1A**). In total, we generated 9 *DAXX* mutant (*DAXX*^*Mut*^) clones: CWR22Rv1 *DAXX*^*Mut*^1 – 4 and LAPC-4 *DAXX*^*Mut*^1 – 5. Three of the four CWR22Rv1 clones (*DAXX*^*Mut*^*1, DAXX*^*Mut*^2, and *DAXX*^*Mut*^*4*) were knockout clones with frameshift mutations in all copies, while the CWR22Rv1 *DAXX*^*Mut*^*3* clone had at least one in-frame mutation (**Fig. 1B**). Although CWR22v1 *DAXX*^*Mut*^*3* had at least one in-frame mutation, all CWR22Rv1 mutant clones had undetectable DAXX protein expression by western blotting (**Fig. 1C**). In contrast, only the LAPC-4 *DAXX*^*Mut*^1 with only frameshift mutations in *DAXX* had undetectable protein, while LAPC-4 *DAXX*^*Mut*^*2, DAXX*^*Mut*^*3, DAXX*^*Mut*^*4*, and *DAXX*^*Mut*^*5* had detectable, although significantly reduced, protein levels (**Fig D**). These observations were consistent with the sequencing results, as LAPC-4 *DAXX*^*Mut*^*2, DAXX*^*Mut*^*3, DAXX*^*Mut*^*4*, and *DAXX*^*Mut*^*5* clones had at least one in-frame mutation in *DAXX* **(Fig. 1B)**.

**Figure 1.**
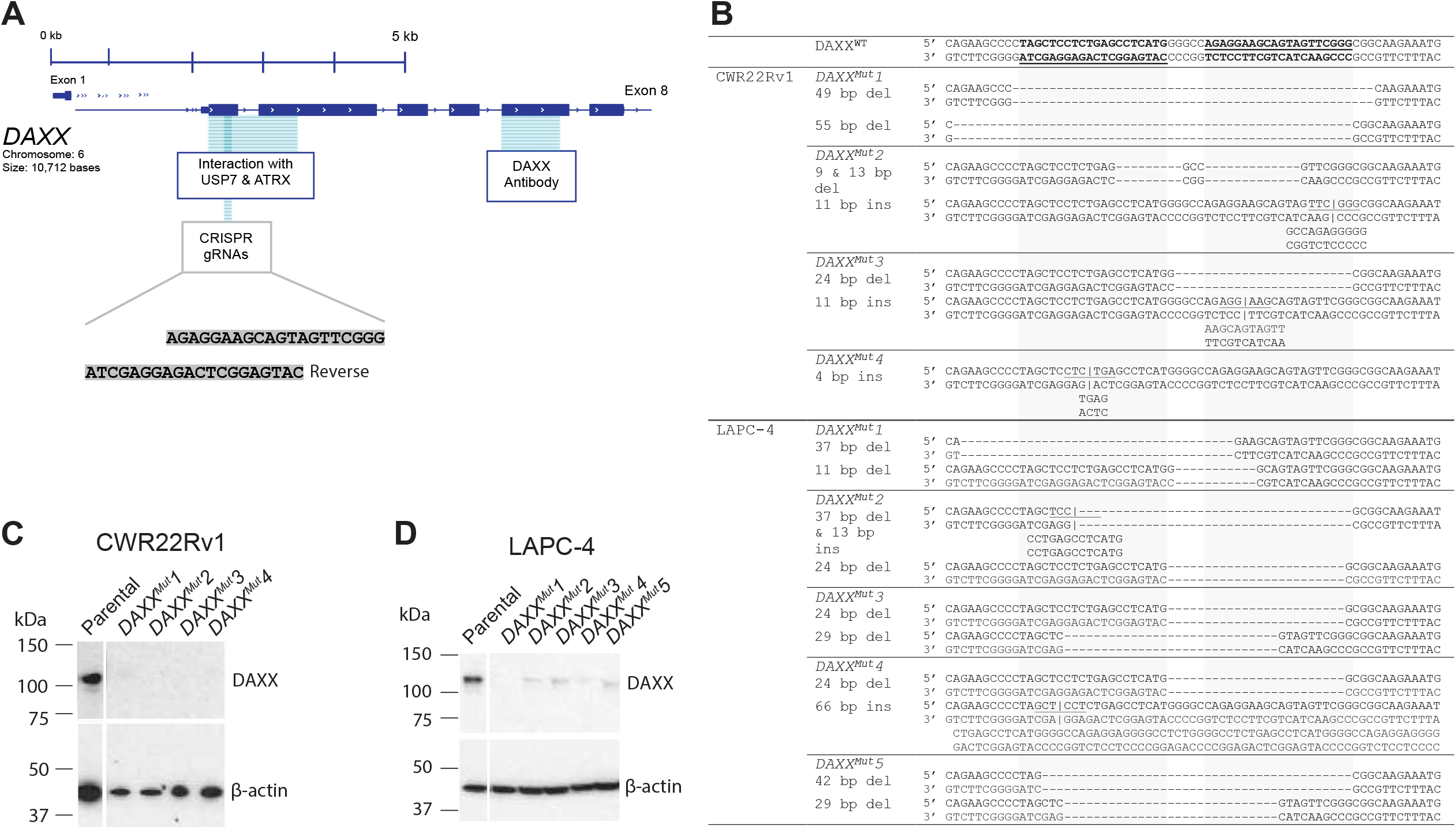
Introducing *DAXX* mutations in CWR22Rv1 and LAPC-4. **(A)** Schematic illustration showing the generation of mutant clones using the CRISPR-Cas9 nickase system, ATRX binding domain, and DAXX antibody binding site. **(B)** Sanger sequencing of CWR22Rv1 and LAPC-4 parental and mutant DAXX clones in the region targeted for disruption by CRISPR. **(C)** Western blot analysis showing loss of DAXX protein expression in all CWR22Rv2 clones, including the clone (CWR22Rv1 *DAXX*^*Mut*^*3*) with at least one in-frame mutation. **(D)** In contrast, all LAPC-4 clones with at least one in-frame mutation in *DAXX* had detectable protein by western blot.

To determine the effect of *DAXX* mutations (including in-frame mutants and knockouts) in CWR22Rv1 and LAPC-4, we assessed the *DAXX*^*Mut*^ clones for ALT-associated hallmarks, including the presence of APBs, C-circles, telomerase activity, and telomere length heterogeneity.

### *DAXX* mutations induce APBs in a subset of clones

ALT-positive cancers have ALT-associated PML bodies (APBs), which are unique nuclear structures that contain donut-shaped promyelocytic leukemia (PML) protein bodies that colocalize with telomeric DNA, the shelterin components TRF1 and TRF2, and proteins involved in DNA synthesis and recombination (12, 13). A telomere FISH assay was performed on fixed cells, followed by immunofluorescence staining for PML bodies to determine the effect of *DAXX* mutations on APB induction. Cancers are generally considered ALT-positive if the ALT-associated ultrabright telomeric foci are present in more than 1% of the cancer cells [24 45 46]. In this study, clones were deemed to possess APB features if the colocalization of the ultrabright telomeric foci and PML signal (**Fig. 2A, Suppl. Fig. 1**) was present in more than 1% of at least 1,000 cells counted. Of the three CWR22Rv1 knockout clones (*DAXX*^*Mut*^*1, DAXX*^*Mut*^*2*, and *DAXX*^*Mut*^*4*) only CWR22Rv1 *DAXX*^*Mut*^4 displayed a significantly higher (>13 fold higher; p< 0.01) percentage of ABP-positive cells (5.15%) compared to the 0.39% in the parental cells (**Fig. 2B**). The percentage of APB-positive cells for CWR22Rv1 *DAXX*^*Mut*^*1, 2*, and *3* cells was comparable to the parental cells, ranging from 0.16% to 0.50%. None of the LAPC4 clones displayed APBs, with the percentage of APB-positive cells ranging from 0% to 0.30% compared to 0.15% in the parental cells (**Fig. 2C**).

**Figure 2.**
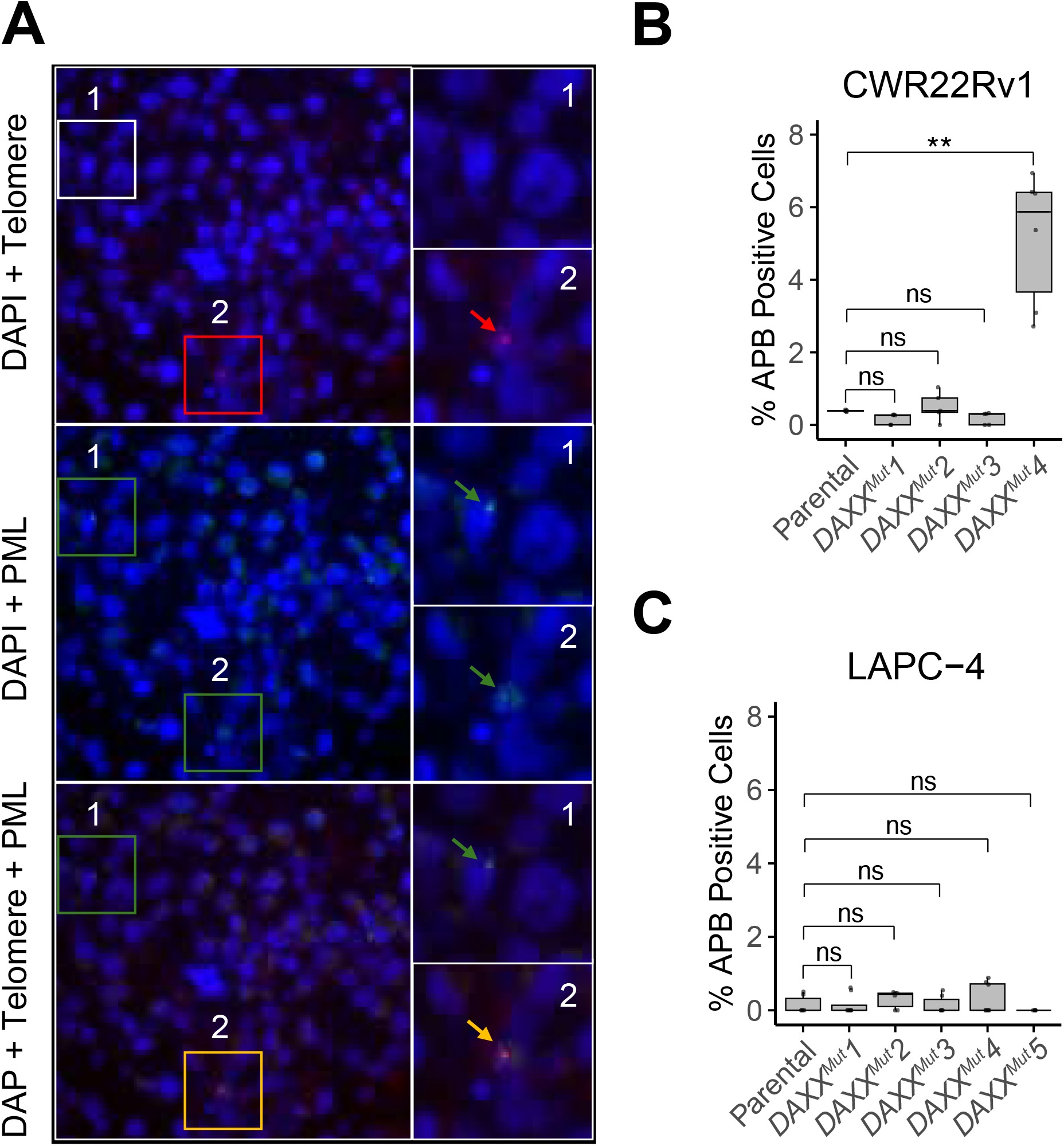
*DAXX* frameshift mutations lead to APB formation in a subset of clones. **(A)** Representative images of APB-positive cells in CWR22Rv1 *DAXX*^*Mut*^*3* clones, indicated by colocalization of Telomere FISH (red arrows/boxes) and PML immunolabeling (green arrows/boxes). APB-positive foci are shown as yellow arrows/boxes in merged images (row 3: DAPI + Telomere + PML). An example of a PML-positive but Telomere-negative cell (APB-negative) is shown in the white box in the DAPI + Telomere panel. Telomere-positive but PML-negative cells were also considered APB-negative. **(B)** Quantification of APB-positive cells in CWR22Rv1 parental and DAXX mutant clones. **(C)** Quantification of APB-positive cells in LAPC-4 parental and DAXX mutant clones.

### *DAXX* mutations induce C-circles in a subset of clones

In general, ALT-positive cells that display APBs will also generate C-circles. These extrachromosomal circular DNAs are single-stranded C-rich telomeric sequences with a segment binding to the complementary G-rich telomeric sequence [24 37]. C-circles are thought to be involved in the ALT process [24 37] and can be detected and quantified using the C-circle assay in which the partial G-strand ([TTAGGG]_n_) segment of the C-circle primes the ϕ29 DNA polymerase. This highly processive DNA polymerase generates telomeric DNA products ≥70 kb [37]. We assessed C-circles in the parental and mutant clones relative to the U2OS cell line (**Fig. 3A**), a well-established ALT-positive cell line that generates C-circles and has no telomerase activity [47]. PC3, an ALT-negative prostate cancer cell line, was used as a negative control. Of all the CWR22Rv1 mutant clones, only CWR22Rv1 *DAXX*^*Mut*^4 had a significant C-circle percentage increase (∼16%) relative to U2OS (**Fig. 3B**). CWR22Rv1 *DAXX*^*Mut*^1, *DAXX*^*Mut*^2, and *DAXX*^*Mut*^3 displayed low levels of C-circle (∼1%) relative to U2OS, which were comparable to the parental CWR22Rv1 (0.98% relative to U2OS). For the LAPC4, the mutant clones exhibited low C-circle levels (<3.5%) relative to those in U2OS (**Fig. 3C**). However, the C-circle levels in some of the clones were significantly higher compared to the parental. The percentage C-circle increases in *DAXX*^*Mut*^1 and *DAXX*^*Mut*^3, relative to U2OS, were 3.6% and 2.2% compared to the <0.1% in the parental.

**Figure 3.**
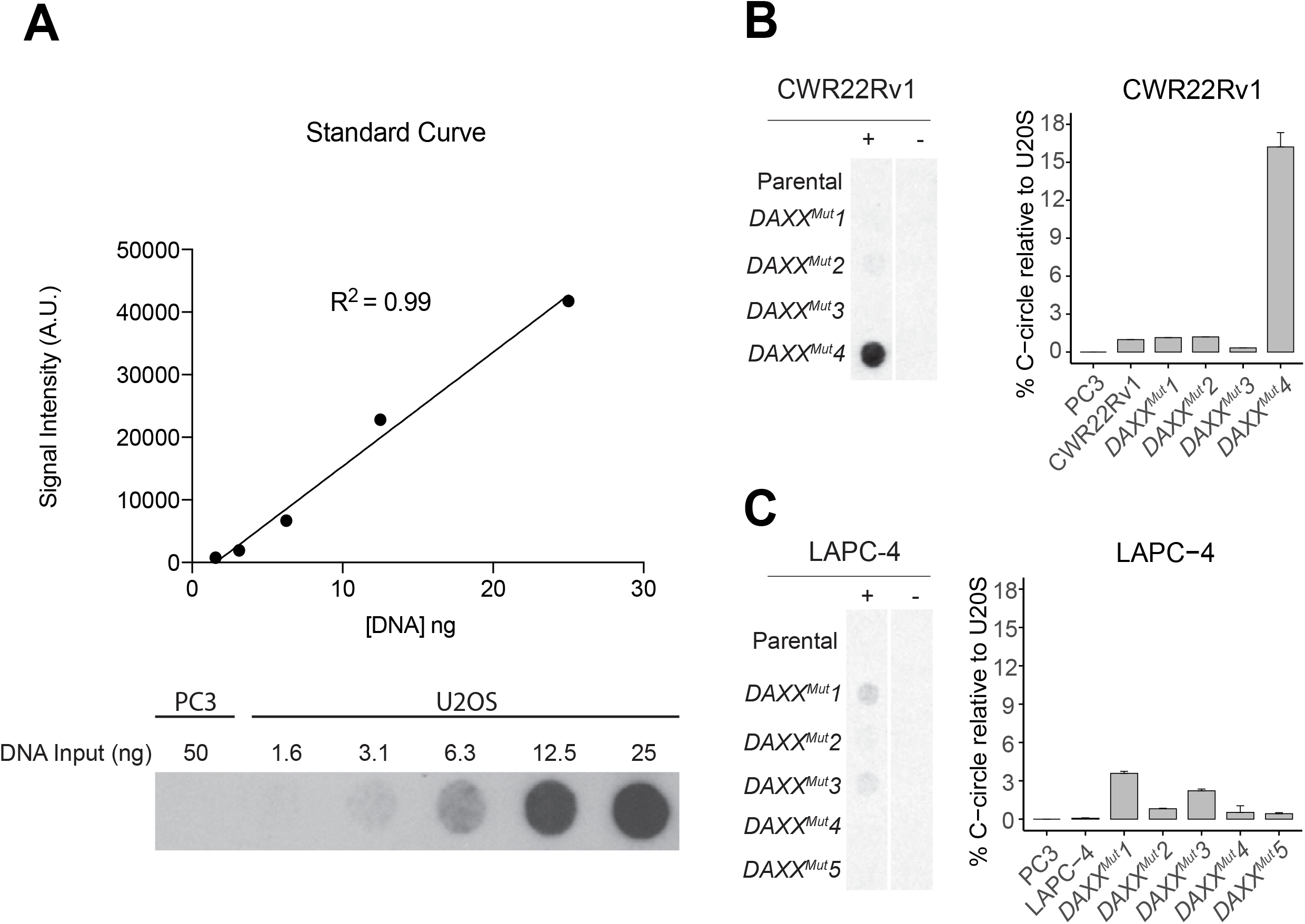
Assessment of C-circle levels of CWR22Rv1 *DAXX*^*Mut*^ and LAPC-4 *DAXX*^*Mut*^ clones. **(A)** A representative standard curve (signal intensity vs DNA input) of detected Ccircles for U2OS, a well-characterized ALT-positive cell line. Below is the representative image of C-circle blots used to plot the standard curves with PC3, an ALT-negative prostate cancer cell line, included as a negative control. Representative C-circle dot blots and the interpolated C-circle levels relative to U2OS of **(B)** CWR22Rv1 parental and mutant clones and **(C)** LAPC-4 parental and mutant clones.

### *DAXX* mutations induce modest telomere length heterogeneity in a subset of clones

Unlike telomerase-expressing cancer cell lines, which exhibit a discernible mean telomere length with telomeres typically ranging a few (<8) kilobases from the mean value, ALT-positive cells display a broad and extremely heterogeneous distribution of telomere lengths ranging from very short (∼10) to greater than 50 kb [48]. We analyzed telomere length heterogeneity using the TRF assay, which probes (TTAGGG)_3_ fragments. As shown by the TRF southern blots in **Fig. 4A** and **B**, except for CWR22Rv1 *DAXX*^*Mut*^*4*, which exhibited modest telomere heterogeneity, none of the clones exhibited increased telomere length and telomere length heterogeneity relative to the parental cell. Interestingly, CWR22Rv1 *DAXX*^*Mut*^3 had shorter overall telomere lengths compared to the parental cells, which can be attributed to background clonal heterogeneity in the parental cell line (**Suppl. Fig. 2**). Additionally, all the clones maintained telomerase activity as shown by TRAP analysis (**Fig. 4C** and **D**), suggesting that *DAXX* mutations did not disrupt telomerase activity, including the CWR22Rv1 *DAXX*^*Mut*^4, which developed some ALT-associated features

**Figure 4.**
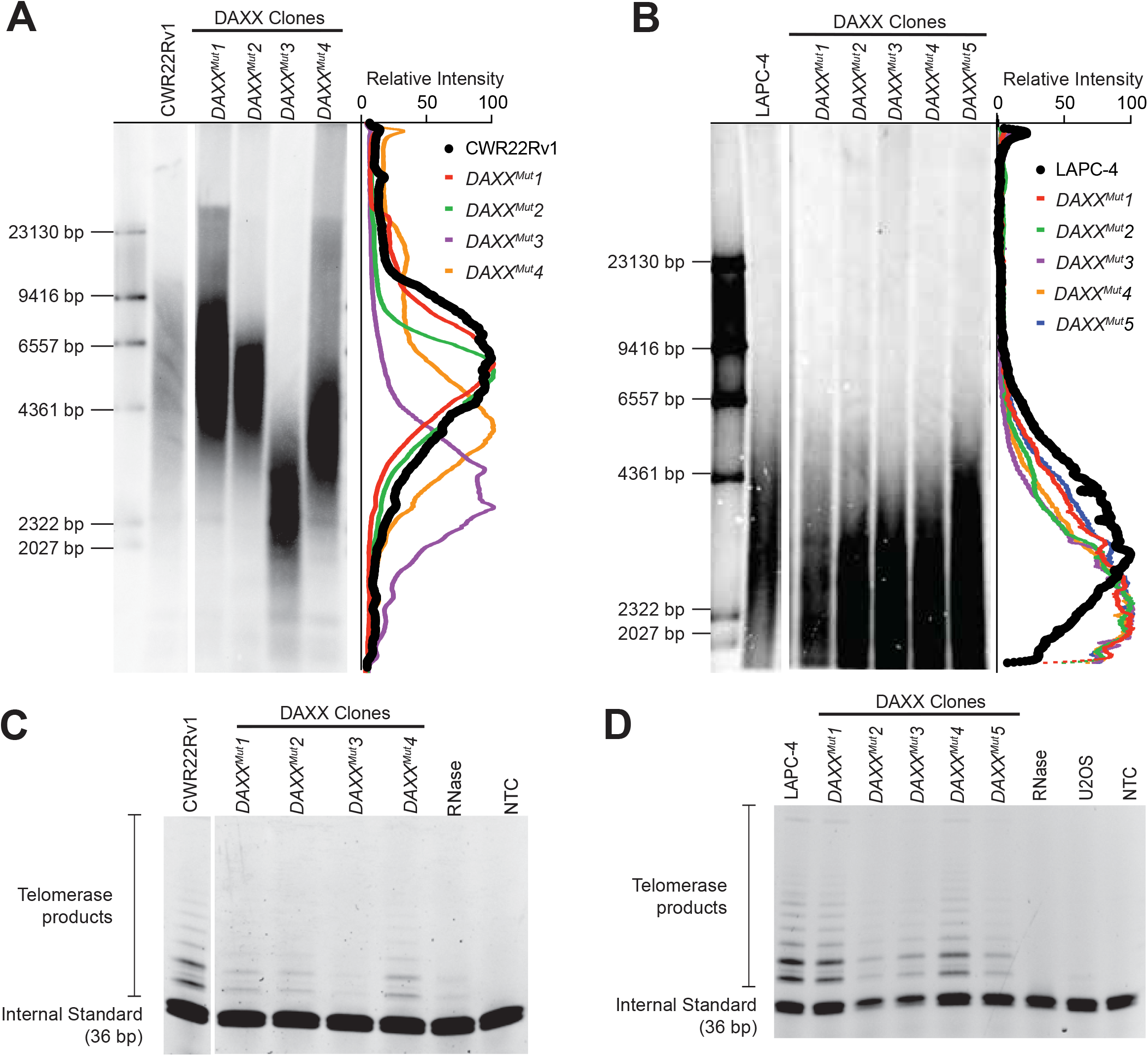
Assessment of telomere lengths and telomerase activity in *DAXX*^*Mut*^ clones. **(A)** and **(B)** Telomere restriction fragment (TRF) southern blots of **(A)** CWR22Rv1 *DAXX*^*Mut*^ and **(B)** LAPC-4 *DAXX*^*Mut*^. Intensity traces of the distribution of TRFs are shown. **(C)** and **(D)** Telomere repeated amplification protocol (TRAP) assay blots of **(C)** CWR22Rv1 *DAXX*^*Mut*^ and **(D)** LAPC-4 *DAXX*^*Mut*^. Telomerase enzymatic activity is detected by visualizing a ladder of template extension products. Negative controls include RNAse-treated cell extract (degrades the telomerase’s RNA telomere repeat template), U2OS (ALT+ human cell line), and a no-template control (NTC).

## DISCUSSION

DAXX and ATRX proteins are binding partners in chromatin remodeling [49 50], and a robust correlation has been previously observed between the presence of mutations in either *DAXX* or *ATRX* and several cancers that display ALT, including PanNETs, pediatric glioblastoma, and uterine leiomyosarcomas [17-19]. In a previous *in-vitro* study that used the same two prostate cancer cell lines used here, we demonstrated that the introduction of *ATRX* loss-of-function mutations induced ALT-associated features in LAPC-4 but not in CWR22Rv1 [24]. Because ATRX and DAXX are binding partners, and mutations in either protein are tightly linked to ALT telomere maintenance programs, we were interested in studying whether mutations in DAXX would also induce ALT-associated hallmarks in LAPC-4.

In the present study, we targeted a sequence in exon 2 that encodes part of the domain of DAXX that interacts with ATRX, forming the ATRX-DAXX complex; the complex plays a vital role in chromatin maintenance by preventing the accumulation of double-strand breaks at telomeric DNA [34 51]. Therefore, we hypothesized that either the loss of DAXX or mutations in exon 2 of DAXX that disrupt the formation of the ATRX-DAXX complex would result in the accumulation of double-strand breaks at telomeric DNA that drives ALT development [52]. We generated four DAXX mutants (CWR22Rv1 *DAXX*^*Mut*^*1, DAXX*^*Mut*^2, and *DAXX*^*Mut*^*4*; and LAPC-4 *DAXX*^*Mut*^1) with frameshift mutations (knockout clones) in *DAXX*, four mutants with both in-frame and frameshift mutations in *DAXX* (CWR22Rv1 *DAXX*^*Mut*^*3* and LAPC-4 *DAXX*^*Mut*^*2, DAXX*^*Mut*^3, and *DAXX*^*Mut*^*5*), and one mutant with only in-frame mutations (LAPC-4 *DAXX*^*Mut*^*4*). Protein expression was not detectable in all the CWR22Rv1 clones, while only one LAPC-4 clone (LAPC-4 *DAXX*^*Mut*^*1*) with frameshift mutations had undetectable protein expression.

Since ATRX loss in LAPC-4 resulted in ALT-associated phenotypes, but did not do so in CWR22Rv1 [24], we expected that loss of DAXX in LAPC-4 would also induce ALT-associated hallmarks. However, we did not see ALT-associated features in any of the LAPC-4 mutant clones, including the LAPC-4 knockout clone (LAPC-4 *DAXX*^*Mut*^1). Interestingly, CWR22Rv1 *DAXX*^*Mut*^*4* (knockout) developed ALT-associated features, including APBs, C-circles, and measurable telomere length heterogeneity. Although the lack of telomerase activity is an ALT hallmark, CWR22Rv1 *DAXX*^*Mut*^*4* maintained its telomerase activity. The present findings suggest that other underlying molecular alterations, in addition to loss of ATRX or DAXX binding, contribute to promoting an ALT-associated phenotype. This would be consistent with differences in how DAXX and ATRX loss impacted the two different prostate cancer cell lines (CWR22rv1 and LAPC-4) and the different clones within each cell line. The findings also align with studies in other cancer types that demonstrate a context-dependent association between the loss of DAXX and/or ATRX expression and ALT-associated features. In PanNETs, inactivating mutations in either the *DAXX* or *ATRX* gene were linked to ALT status [45]. Similarly, in uterine leiomyosarcomas, inactivating mutations in either *ATRX* or *DAXX* were associated with ALT [19]. Likewise, in cancers of the central nervous system, only mutations in *ATRX*, not *DAXX*, were associated with ALT; in fact, *DAXX* gene mutations were undetectable in the studied cohort [45]. Similarly, in pediatric glioblastoma, ATRX protein loss was strongly correlated with ALT, although some ATRX-positive cases were also ALT-positive [18]. In an *in-vitro* study of human glioma cell lines, ATRX loss induced hallmarks of ALT in some cell lines, although telomerase activity and overall telomere length heterogeneity remain unaffected [53]. Moreover, loss of ATRX protein and mutations in the *ATRX* gene are hallmarks of ALT-immortalized cell lines [54].

Here, we evaluated the effect of loss of DAXX expression on ALT-associated features in prostate cancer cell lines. Together with our previous ATRX knockout models [24], the models generated in this study provide useful molecular tools. Of particular interest, future experiments examining whether the loss of telomerase or inhibition of telomerase activity is sufficient to convert these DAXX mutants to ALT-dependent cell lines will provide additional insights into the role of DAXX and telomere maintenance. Beyond understanding ALT biology, these models will be useful for probing DAXX-related biology, including its role in prostate carcinogenesis.

## Acknowledgements

This work was supported by National Institutes of Health (5T32CA009110-38 to M.K.G., F32CA213742 to M.K.G., 5R01CA172380-05 to A.K.M., P30 CA006973 to the Sidney Kimmel Comprehensive Cancer Center); American Cancer Society (Boston University-Boston Medical Center Pilot and Feasibility Program to C.M. Heaphy). We wish to thank the staff at the Flow Cytometry and Cell Sorting Core Facility at Johns Hopkins School of Public Health, and the Oncology Tissue Core and Microarray Core at Johns Hopkins Sidney Kimmel Comprehensive Cancer Center.

## Conflicts of Interest

There are no conflicts of interest to declare.

## Notes

### Competing Interest Statement

The authors have declared no competing interest.

## REFERENCES

1. O’Sullivan RJ, Karlseder J. Telomeres: protecting chromosomes against genome instability. Nat Rev Mol Cell Biol 2010;11(3):171–81

2. Zvereva MI, Shcherbakova DM, Dontsova OA. Telomerase: structure, functions, and activity regulation. Biochemistry (Mosc) 2010;75(13):1563–83

3. Thomson JA, Itskovitz-Eldor J, Shapiro SS, et al. Embryonic stem cell lines derived from human blastocysts. Science 1998;282(5391):1145–7

4. Hiyama E, Hiyama K. Telomere and telomerase in stem cells. Br J Cancer 2007;96(7):1020–4

5. Shay JW, Bacchetti S. A survey of telomerase activity in human cancer. Eur J Cancer 1997;33(5):787–91

6. Olovnikov AM. A theory of marginotomy. The incomplete copying of template margin in enzymic synthesis of polynucleotides and biological significance of the phenomenon. J Theor Biol 1973;41(1):181–90

7. Vaziri H. Critical telomere shortening regulated by the ataxia-telangiectasia gene acts as a DNA damage signal leading to activation of p53 protein and limited life-span of human diploid fibroblasts. A review. Biochemistry (Mosc) 1997;62(11):1306–10

8. Dagg RA, Pickett HA, Neumann AA, et al. Extensive Proliferation of Human Cancer Cells with Ever-Shorter Telomeres. Cell Rep 2017;19(12):2544–56

9. Viceconte N, Dheur MS, Majerova E, et al. Highly Aggressive Metastatic Melanoma Cells Unable to Maintain Telomere Length. Cell Rep 2017;19(12):2529–43

10. Hanahan D, Weinberg RA. Hallmarks of cancer: the next generation. Cell 2011;144(5):646–74

11. Shay JW, Reddel RR, Wright WE. Cancer. Cancer and telomeres--an ALTernative to telomerase. Science 2012;336(6087):1388–90

12. Bryan TM, Englezou A, Dalla-Pozza L, Dunham MA, Reddel RR. Evidence for an alternative mechanism for maintaining telomere length in human tumors and tumor-derived cell lines. Nat Med 1997;3(11):1271–4

13. Dilley RL, Verma P, Cho NW, Winters HD, Wondisford AR, Greenberg RA. Break-induced telomere synthesis underlies alternative telomere maintenance. Nature 2016;539(7627):54–58

14. Zhang JM, Yadav T, Ouyang J, Lan L, Zou L. Alternative Lengthening of Telomeres through Two Distinct Break-Induced Replication Pathways. Cell Rep 2019;26(4):955–68

15. Drane P, Ouararhni K, Depaux A, Shuaib M, Hamiche A. The death-associated protein DAXX is a novel histone chaperone involved in the replication-independent deposition of H3.3. Genes Dev 2010;24(12):1253–65

16. Goldberg AD, Banaszynski LA, Noh KM, et al. Distinct factors control histone variant H3.3 localization at specific genomic regions. Cell 2010;140(5):678–91

17. Heaphy CM, Subhawong AP, Hong SM, et al. Prevalence of the alternative lengthening of telomeres telomere maintenance mechanism in human cancer subtypes. Am J Pathol 2011;179(4):1608–15

18. Schwartzentruber J, Korshunov A, Liu XY, et al. Driver mutations in histone H3.3 and chromatin remodelling genes in paediatric glioblastoma. Nature 2012;482(7384):226–31

19. Makinen N, Aavikko M, Heikkinen T, et al. Exome Sequencing of Uterine Leiomyosarcomas Identifies Frequent Mutations in TP53, ATRX, and MED12. PLoS Genet 2016;12(2):e1005850

20. Lindsay CR, Giovinazzi S, Ishov AM. Daxx is a predominately nuclear protein that does not translocate to the cytoplasm in response to cell stress. Cell Cycle 2009;8(10):1544–51

21. Song JJ, Lee YJ. Role of the ASK1-SEK1-JNK1-HIPK1 signal in Daxx trafficking and ASK1 oligomerization. J Biol Chem 2003;278(47):47245–52

22. Jia L, Yu W, Wang P, Li J, Sanders BG, Kline K. Critical roles for JNK, c-Jun, and Fas/FasL-Signaling in vitamin E analog-induced apoptosis in human prostate cancer cells. Prostate 2008;68(4):427–41

23. Pinto LM, Pailas A, Bondarchenko M, et al. DAXX promotes centromeric stability independently of ATRX by preventing the accumulation of R-loop-induced DNA double-stranded breaks. Nucleic Acids Res 2024;52(3):1136–55

24. Graham MK, Kim J, Da J, et al. Functional Loss of ATRX and TERC Activates Alternative Lengthening of Telomeres (ALT) in LAPC4 Prostate Cancer Cells. Mol Cancer Res 2019;17(12):2480–91

25. Lin DY, Fang HI, Ma AH, et al. Negative modulation of androgen receptor transcriptional activity by Daxx. Mol Cell Biol 2004;24(24):10529–41

26. Song MS, Salmena L, Carracedo A, et al. The deubiquitinylation and localization of PTEN are regulated by a HAUSP-PML network. Nature 2008;455(7214):813–7

27. Jamaspishvili T, Berman DM, Ross AE, et al. Clinical implications of PTEN loss in prostate cancer. Nat Rev Urol 2018;15(4):222–34

28. Tsourlakis MC, Schoop M, Plass C, et al. Overexpression of the chromatin remodeler death-domain-associated protein in prostate cancer is an independent predictor of early prostate-specific antigen recurrence. Hum Pathol 2013;44(9):1789–96

29. Kwan PS, Lau CC, Chiu YT, et al. Daxx regulates mitotic progression and prostate cancer predisposition. Carcinogenesis 2013;34(4):750–9

30. Puto LA, Brognard J, Hunter T. Transcriptional Repressor DAXX Promotes Prostate Cancer Tumorigenicity via Suppression of Autophagy. J Biol Chem 2015;290(25):15406–20

31. Puto LA, Benner C, Hunter T. The DAXX co-repressor is directly recruited to active regulatory elements genome-wide to regulate autophagy programs in a model of human prostate cancer. Oncoscience 2015;2(4):362–72

32. Sramkoski RM, Pretlow TG, 2nd, Giaconia JM, et al. A new human prostate carcinoma cell line, 22Rv1. In Vitro Cell Dev Biol Anim 1999;35(7):403–9

33. Klein KA, Reiter RE, Redula J, et al. Progression of metastatic human prostate cancer to androgen independence in immunodeficient SCID mice. Nat Med 1997;3(4):402–8

34. Mahmud I, Liao D. DAXX in cancer: phenomena, processes, mechanisms and regulation. Nucleic Acids Res 2019;47(15):7734–52

35. Hsu PD, Scott DA, Weinstein JA, et al. DNA targeting specificity of RNA-guided Cas9 nucleases. Nat Biotechnol 2013;31(9):827–32

36. Schindelin J, Arganda-Carreras I, Frise E, et al. Fiji: an open-source platform for biological-image analysis. Nat Methods 2012;9(7):676–82

37. Henson JD, Cao Y, Huschtscha LI, et al. DNA C-circles are specific and quantifiable markers of alternative-lengthening-of-telomeres activity. Nat Biotechnol 2009;27(12):1181–5 doi: 10.1038/nbt.1587 [published Online First: Epub Date]|.

38. Mender I, Shay JW. Telomerase Repeated Amplification Protocol (TRAP). Bio Protoc 2015;5(22)

39. Kimura M, Stone RC, Hunt SC, et al. Measurement of telomere length by the Southern blot analysis of terminal restriction fragment lengths. Nat Protoc 2010;5(9):1596–607

40. Lee H, Birren B, Lai E. Ultraviolet nicking of large DNA molecules from pulsed-field gels for southern transfer and hybridization. Anal Biochem 1991;199(1):29–34

41. Ran FA, Hsu PD, Wright J, Agarwala V, Scott DA, Zhang F. Genome engineering using the CRISPR-Cas9 system. Nat Protoc 2013;8(11):2281–308

42. Tang J, Wu S, Liu H, et al. A novel transcription regulatory complex containing death domain-associated protein and the ATR-X syndrome protein. J Biol Chem 2004;279(19):20369–77

43. Wang X, Zhao Y, Zhang J, Chen Y. Structural basis for DAXX interaction with ATRX. Protein Cell 2017;8(10):767–71

44. Giovinazzi S, Morozov VM, Summers MK, Reinhold WC, Ishov AM. USP7 and Daxx regulate mitosis progression and taxane sensitivity by affecting stability of Aurora-A kinase. Cell Death Differ 2013;20(5):721–31

45. Heaphy CM, de Wilde RF, Jiao Y, et al. Altered telomeres in tumors with ATRX and DAXX mutations. Science 2011;333(6041):425

46. Haffner MC, Mosbruger T, Esopi DM, et al. Tracking the clonal origin of lethal prostate cancer. J Clin Invest 2013;123(11):4918–22

47. Scheel C, Schaefer KL, Jauch A, et al. Alternative lengthening of telomeres is associated with chromosomal instability in osteosarcomas. Oncogene 2001;20(29):3835–44

48. Bryan TM, Englezou A, Gupta J, Bacchetti S, Reddel RR. Telomere elongation in immortal human cells without detectable telomerase activity. EMBO J 1995;14(17):4240–8

49. Lewis PW, Elsaesser SJ, Noh KM, Stadler SC, Allis CD. Daxx is an H3.3-specific histone chaperone and cooperates with ATRX in replication-independent chromatin assembly at telomeres. Proc Natl Acad Sci U S A 2010;107(32):14075–80

50. Xue Y, Gibbons R, Yan Z, et al. The ATRX syndrome protein forms a chromatin-remodeling complex with Daxx and localizes in promyelocytic leukemia nuclear bodies. Proc Natl Acad Sci U S A 2003;100(19):10635–40

51. Hoelper D, Huang H, Jain AY, Patel DJ, Lewis PW. Structural and mechanistic insights into ATRX-dependent and -independent functions of the histone chaperone DAXX. Nat Commun 2017;8(1):1193

52. Cho NW, Dilley RL, Lampson MA, Greenberg RA. Interchromosomal homology searches drive directional ALT telomere movement and synapsis. Cell 2014;159(1):108–21

53. Brosnan-Cashman JA, Yuan M, Graham MK, et al. ATRX loss induces multiple hallmarks of the alternative lengthening of telomeres (ALT) phenotype in human glioma cell lines in a cell line-specific manner. PLoS One 2018;13(9):e0204159

54. Lovejoy CA, Li W, Reisenweber S, et al. Loss of ATRX, genome instability, and an altered DNA damage response are hallmarks of the alternative lengthening of telomeres pathway. PLoS Genet 2012;8(7):e1002772

