## Supplemental Figure 1 for "Loss of DAXX induces alternative lengthening of telomeres (ALT)-associated hallmarks in prostate cancer cells in a context-specific manner"

Supplementary Figure 1. Representative images of Telomere (top) and PML (middle) staining in LAPC-4 clones. The LAPC-4 clones were mostly APB negative, shown by the absence of telomere signal, hence the absence of PML and Telomere colocalization.

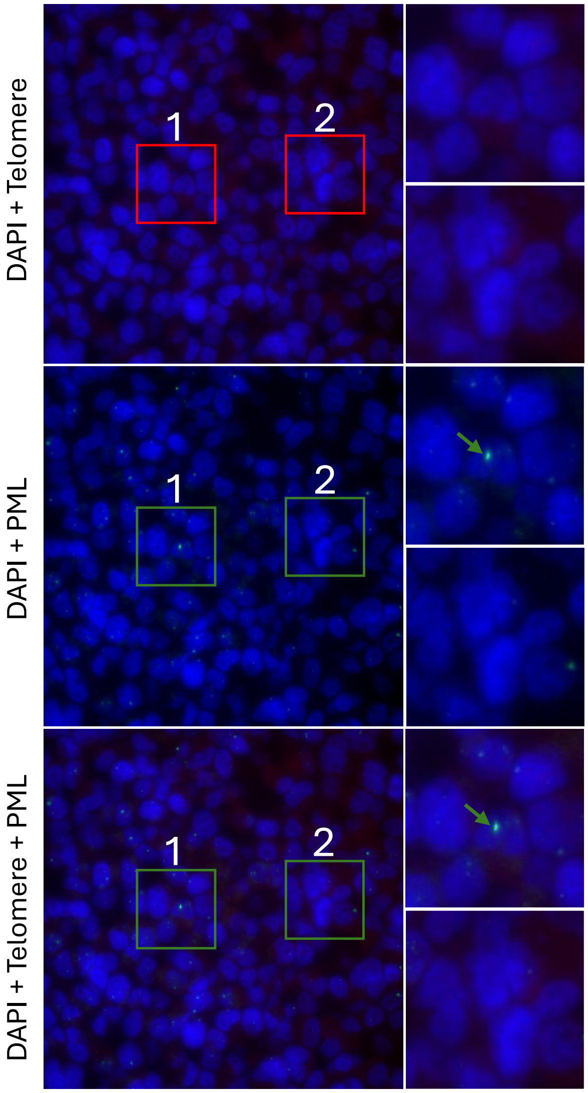
