## Supplemental Figure 2 for "Loss of DAXX induces alternative lengthening of telomeres (ALT)-associated hallmarks in prostate cancer cells in a context-specific manner"

Supplementary Figure 2. Telomere length background clonal heterogeneity in the parental cell line

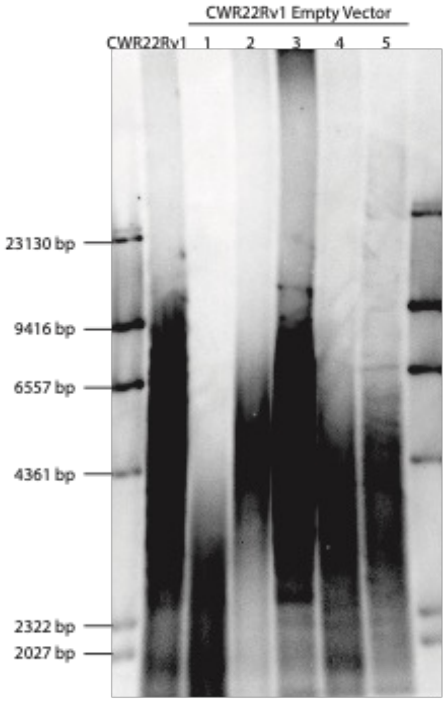


| Cell Line | Size (bp) |
|---|---|
| CWR22Rv1 | 3072 |
| CWR22Rv1 EV 1 | 2302 |
| CWR22Rv1 EV 2 | 4599 |
| CWR22Rv1 EV 3 | 4982 |
| CWR22Rv1 EV 4 | 3610 |
| CWR22Rv1 EV 5 | 3680 |

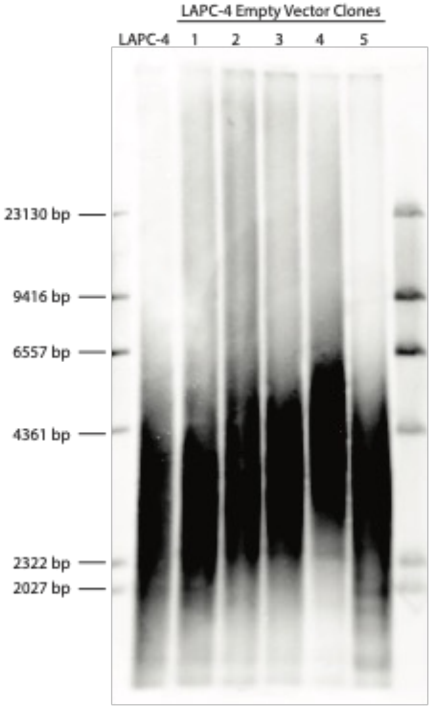


| Cell Line | Size (bp) |
|---|---|
| LAPC-4 | 3072 |
| LAPC-4 EV 1 | 2912 |
| LAPC-4 EV 2 | 3167 |
| LAPC-4 EV 3 | 3466 |
| LAPC-4 EV 4 | 4225 |
| LAPC-4 EV 5 | 3347 |
